# panX: pan-genome analysis and exploration

**DOI:** 10.1101/072082

**Authors:** Wei Ding, Franz Baumdicker, Richard A. Neher

## Abstract

Horizontal transfer, gene loss, and duplication result in dynamic bacterial genomes shaped by a complex mixture of different modes of evolution. Closely related strains can differ in the presence or absence of many genes, and the total number of distinct genes found in a set of related isolates – the pan-genome – is often many times larger than the genome of individual isolates. We have developed a pipeline that efficiently identifies orthologous gene clusters in the pan-genome. This pipeline is coupled to a powerful yet easy-to-use web-based visualization software for interactive exploration of the pan-genome. The visualization consists of connected components that allow rapid filtering and searching of genes and inspection of their evolutionary history. For each gene cluster, panX displays an alignment, a phylogenetic tree, maps mutations within that cluster to the branches of the tree and infers gain and loss of genes on the core-genome phylogeny. PanX is available at pangenome.de. Custom pan-genomes can be visualized either using a webserver or by serving panX locally as a browser-based application.

## Introduction

In addition to vertically passing down their genome to offspring, bacteria have the capability to acquire genetic material from the environment via horizontal transfer [1]. Genes are transferred among bacteria by a variety of mechanisms including active uptake, mobile genetic elements, and gene transfer by viruses [2]. In addition to gene gain, genes are frequently duplicated or lost. The mix of vertical transmission and horizontal transfer complicates the phylogenetic analysis of bacterial genomes and results in patterns of genetic diversity that are difficult to interpret [3].

A common approach when analyzing collections of bacterial genomes is categorizing genes into the *core* or *accessory* genome [4–6]. Core genes are shared by all strains in a group of isolates, accessory genes shared by two or more but not all strains, and unique genes are specific to a single strain. The union of all genes found in a group of strains (e.g. strains from one species) is called the pan-genome, which is typically several times larger than the core genome. The core genome is often used to assess the relatedness among the genomes in the sample and to approximate the species tree, but extensive horizontal transfer has been documented in the core genome as well [7] such a tree reconstructed from core genome diversity does not necessarily reflect the phylogeny. Different software tools try to infer or remove the impact of recombination on the species level phylogeny [8, 9].

By providing a repertoire of functional genes, gene gain from the pan-genome can facilitate the acquisition of new metabolic pathways [10], the adaptation to new habitats, or the emergence of drug resistant variants [11]. With the rapidly increasing number of sequenced bacterial genomes, it is now possible to detect associations between metadata such as habitats, phenotypes, clinical manifestations and the presence or absence of particular genes [12, 13].

Pan-genome construction from a group of related bacterial genomes typically involves the identification of homologous regions by all-against-all comparisons followed by clustering orthologous genes [4]. Several software packages and pipelines have been developed to construct such pan-genomes that differ in the heuristics used to compare strains and generate clusters [14–17].

One fundamental limitation, however, is the difficulty to interrogate, explore, and visualize the pan-genome and the evolutionary relationships between strains. In absence of recombination, the purely vertical evolutionary history of strains would be represented by a single phylogenetic tree, the species tree. With horizontal transfer, the history of different loci in the genome is described by different trees resulting in a phylogenetic forest or network [18, 19]. While phylogenetic networks can be visualized using consensus representations such as split networks [20], the history and distribution of individual proteins are often critical, for example when searching for associations with phenotypes like drug resistance. Individual clusters of orthologous sequences, however, can again be represented by a tree if genes are short enough that recombination within the gene can be ignored. Some gene trees might be similar to the species tree, while others might deviate dramatically from the species tree. The degree of incongruence of the gene tree with the species tree contains important information about the dynamics of gene gain and loss.

Here, we present panX, a web-based environment for microbial pan-genome data visualization and exploration based on an automated pan-genome identification pipeline. The pipeline breaks the genomes of a large number of annotated genomes (e.g. NCBI reference sequences) into genes and then clusters genes into orthologous groups. From these clusters, panX identifies the core genome, builds a strain-level phylogeny using SNPs in the core genome, constructs multiple alignments of sequences in gene clusters, builds trees for individual genes and maps the gene presence/absence pattern onto the core genome tree. The interactive browser-based application then allows the exploration of the above features and provides flexible filter, sort, and search functionalities. This application is available at pangenome.de with a collection of pan-genomes prepared by us, but can also be deployed on other servers with custom pan-genomes, or can be run locally as a browser-based desktop application.

## Materials and Methods

### Identification of orthologous gene clusters

The initial steps in the computational pipeline underlying panX (illustrated in Fig. 1) is broadly similar to other tools used to construct pan-genomes [14–17, 24]. PanX algorithm identifies groups of homologous genes by similarity search using DIAMOND and clustering by MCL. In a second step, panX builds phylogenies of these groups of genes and splits them into approximately orthologous clusters by examining the structure of the trees. PanX thus combines the speed of graph methods to identify groups of similar sequences with tree based methods applied to individual clusters to accurately split homologous sequences into orthologous groups.

**FIG. 1:**
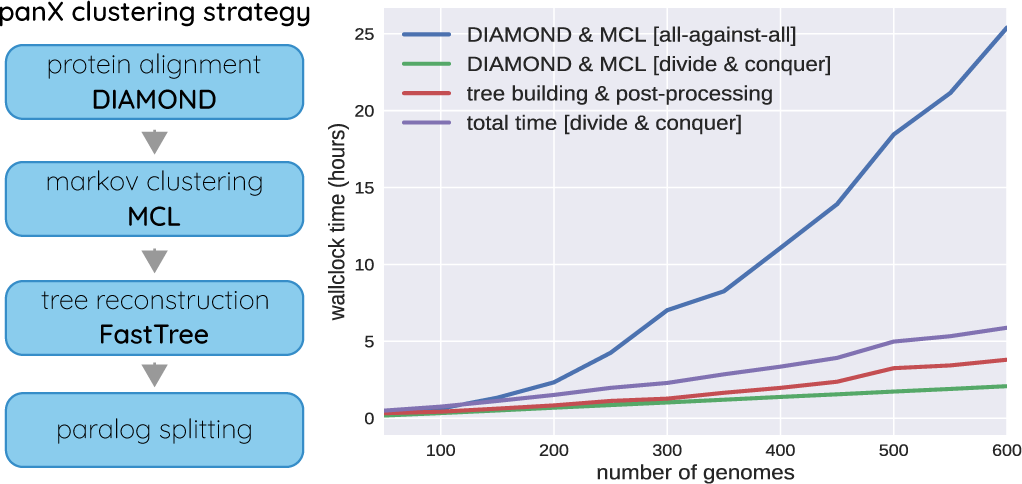
panX analysis pipeline: panX uses DIAMOND [21] and MCL [22, 23] to identify clusters of homologous genes from a collection of annotated genomes. These clusters are then analyzed phylogenetically and split into orthologous groups based on the tree structure. The graph on the right show the time required to identify orthologous clusters in pan-genomes of different size on a compute node with 64 cores. The naive all-against-all comparison with DIAMOND scales quadratically with the number of genomes (blue line, ‘DIAMOND & MCL [all-against-all]). The “divide and conquer” strategy where clustering is first applied to batches of sequences and batches are subsequently clustered (see text) reduces this scaling to approximately linear (green line). Tree building and post-processing take about as long as the clustering itself for pan-genomes of 500 genomes.

#### Identification of groups of homologous sequences

As input, panX uses annotated genome sequences in GenBank format. To identify homologous proteins, panX performs an all-against-all similarity search using DIAMOND [21] with default e-value cut-off of 0.001. From the diamond output, panX constructs a file listing pairs of genes and their bitscore. Using bitscore instead of e-value avoids underflow problems and combines similarity and length of the homologous region [25]. The table of similarity scores serves as input for the Markov Clustering Algorithm (MCL) [22, 23] to create the clusters of putatively orthologous genes. The DIAMOND similarity search can be multi-threaded and panX uses 64 CPUs by default if run on a compute cluster. Since DIAMOND aligns proteins, ribosomal RNAs rRNAs have to be handled separately. PanX extracts rRNAs from GenBank files and compares sequences to each other via blastn. The output of blastn is then processed in the same way as the protein comparison by DIAMOND, that is hits are clustered by MCL and clustered refined using the phylogeny based post-processing (see below). If desired, DIAMOND similarity search can be replaced completely by other sequence similarity search tools such as blastx or blastn.

*Divide-and-conquer strategy for large data set:* The all-against-all similarity search scales quadratically with the number of genomes making the naive implementation infeasible for thousands of genomes, see Fig. 1. However, the majority of these comparisons are redundant and can be avoided by first clustering small batches of genomes and subsequently combining different batches. Specifically, we apply the DIAMOND and MCL steps to subsets of 50 genomes (large enough to benefit from DIAMONDS double indexing strategy, small enough such that the all-against-all is not yet prohibitive) and derive gene clusters of this “sub-pan-genome”. Each gene cluster is then reduced to a representative sequence and the representative sequences of all gene clusters are used as a “pseudo genome” representing the entire batch. The pseudo genomes representing the different batches are then again clustered using the DIAMOND+MCL steps. Eventually, complete clusters are constructed by combining sequences represented by the pseudo genomes. This “divideand-conquer” strategy can be applied repeatedly for very large pan-genomes and keeps scaling of clustering approximately linear, see Fig. 1.

#### Splitting into orthologous clusters

In our experience, it is advisable to cluster proteins aggressively and split clusters with paralogous sequences in a post-processing step. The groups of paralogs are often readily apparent in a phylogenetic tree. PanX reconstructs trees from sequences in each cluster by first aligning the protein sequences using MAFFT [26]. The protein alignment is then used to construct a codon-alignment of the corresponding nucleotide sequences by inserting a gap of length three for every gap in the amino acid alignment. From nucleotide sequence alignment panX then reconstructs a tree using FastTree [27]. The runtime of FastTree scales approximately as *n*^3/2^ with the number of sequences. While superlinear, this scaling still allows the analysis of thousands of genomes. For 600 genomes, tree building and post-processing takes about twice as long as the initial clustering (see Fig. 1). Once the tree of a cluster is available, panX employs a three-step procedure to decide whether and where a cluster should be split into sub-clusters.

*Splitting distantly related homologs:* Since branch lengths reflect evolutionary distances among genes within one cluster, groups of distantly related genes, for example resulting from an ancient duplication, are connected by long branches and can be easily spotted in a gene tree – at least for pan-genome of low or moderate diversity. PanX splits trees into subtrees at branches whose length exceeds an adaptive threshold. This threshold is determined from the average diversity *d_c_* of single copy core genes via

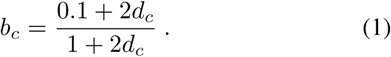

This cut-off increases as 0.1 + 2*d_c_* for very similar strains and eventually saturates at 1. The genetic diversity of the core genes will be of the same order of magnitude as mutational distance to the most recent common ancestor (MRCA) of the collection of genomes. Branches much longer than this diversity will typically correspond to duplications long before the MRCA. Hence, the cluster should be cut along these branches. For very diverse pan-genome with *d_c_ >* 0.25, however, this simple threshold splitting will result in under-clustering and should be switched off.

The newly formed clusters are then re-aligned, a new tree is built, and further split using above-mentioned method until no long branches can be detected.

*Splitting closely related paralogs:* Splitting branches longer than *b_c_* will miss recent gene duplication events. To detect paralogous clusters more sensitively, panX calculates a paralogy score for each branch in each tree. The paralogy score of a branch is the number of strains represented on both sides of the branch in the phylogeny. This score can be calculated in linear time for all branches simultaneously in two tree traversals. Clusters are then split into two sub-clusters if the highest paralogy score *ϕ_max_* and the length ℓ of the corresponding branch fulfill the following criteria:

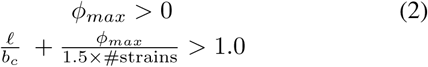

Here *b_c_* refers to the cut-off defined in Eq. (1). The criterion *ϕ_max_ >* 0 prevents splitting irrespective of paralogy. Other criteria could be used but in our experience, this linear discriminator works well for many different applications.

This paralog splitting is iterated until no gene cluster is split. Some heavily duplicated genes require more than five rounds of splitting. The parameters of this splitting step can be set by the user.

*Merging fragmented clusters* A small number of genes are not properly clustered either because no homology was detected initially or the clustering by MCL failed. Such unclustered sequences manifest themselves as many singleton clusters of identical length. To detect those sequences, panX calculates the average length of sequences in each cluster and searches for peaks in the distribution of gene cluster length. Unclustered genes show as spikes in this empirical gene length distribution, which panX can identify by detecting peaks in this distribution relative to a smoothed background distribution. For each detected peak, all involved genes are gathered in one pre-cluster, their sequences are aligned and the corresponding phylogeny is inferred. Sub-clusters are then split following the same long branch splitting principle as described above. Currently, only sequences of identical length are combined into tentative clusters. This condition could be relaxed but this has not been necessary in our experience.

The number of clusters that require post-processing depends on the diversity of the data set. On simulated data, roughly 40% of initial clusters need splitting for the least diverse sets, while only a small fraction of clusters required post-processing for the more diverse data sets.

### Phylogenetic analysis of gene clusters

To reduce the computational burden of the subsequent visualization of the pan-genome, alignments, trees, and other properties of the gene clusters are precomputed. The input for this phylogenetic analysis is either the output of the panX pipeline presented above, or the output of Roary. Other pangenome tools could be used when a script parsing the clustering output is supplied.

*Tree building and ancestral reconstruction.* PanX extracts all variable positions from the nucleotide alignments of all single copy core genes (those gene clusters in which all strains are represented exactly once) to construct a core-genome SNP matrix. This SNP matrix is used to build a core genome phylogenetic tree using FastTree [27], which is further refined by RaxML [28] following a similar strategy as implemented in nextflu [29]. Due to homologous recombination, this core genome tree may not reflect the true history for each of the genes in the core genomes [7] and branch lengths do not reflect sequence similarity since only variable sites are used [30]. Nevertheless, this core genome SNP phylogeny is still a useful approximation of the relationships of the different strains that can be used as a scaffold to investigate the evolution of the mobile genome and the distribution of phenotypes.

Phylogenetic trees for a gene cluster have already been inferred in the cluster post-processing step. PanX uses these trees to infer ancestral sequences of internal nodes using a joint maximum likelihood approach [31] as implemented in TreeTime [**?**]. Likely mutations are mapped onto the branches of the tree using this ancestral reconstruction.

Then, we infer the presence or absence of each gene cluster on internal nodes of the core genome SNP tree using an analogous ancestral inference procedure. Individual gain and loss events are associated with branches based on this ancestral reconstruction. The gain and loss rates are optimized such that the likelihood for the observed presence/absence pattern of genes is maximized [32, 33]. We found that optimal loss rates are always larger than the gain rates but their ratio is variable among species with a median ratio of 22 (inter-quartile range 9 to 35).

Gene clusters, trees, mutations, and metadata are stored as JSON files for the web visualization.

### Associations

If informative numerical meta data are attached to the genomes, panX can quantify the association genetic signatures with meta data. PanX considers associations of two types: Either particular variants of a gene are associated with a phenotype or the presence or absence of a gene can be linked to a phenotype. PanX can calculate an association score for both types of association for each gene cluster.

To quantify how well a phenotype is associated with particular variants of a gene, panX computes a normalized difference of phenotypes of strains on either side of a branch in the tree as follows

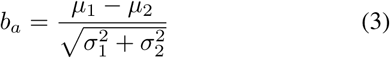

where *µ*_1/2_ and 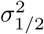 are the mean and variance of the phenotypes of either side of the branch, respectively. This score is calculated for every branch of the tree and the maximum score is reported.

To quantify associations of a phenotype with the presence and absence of a gene, panX uses the average phenotype *µ_p_* of strains carrying the gene and the average *µ_a_* of strains without the gene, the overall variance of the phenotype *σ*^2^, and the number of gain/loss events *n* to calculate the score

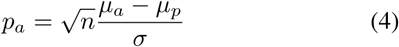

A simple score like the above won’t reliably separate true associations from all false positives but these scores can be very useful to prioritize gene cluster for detailed follow-up investigation. The web application allows to sort genes by their association scores such that strongly associated genes can be rapidly detected and inspected.

### Simulation of pan-genomes

To assess the accuracy of clustering methods we simulated 120 pan-genomes with 30 artificial genomes each. The simulation evolves ancestral sequences along coalescent trees generated by the software ms [34] and allows for horizontal transfer as well as gene loss and gain. To get realistic ancestral sequences we used one representative gene from each KEGG ortholog group [35] present in the *E. coli* strain K-12 (NC 000913) as a starting point for the simulation. This yielded 2803 different genes.

Using these as ancestral sequences, we simulated pangenomes by the following procedure: For each of the 2803 genes we generated correlated trees using the software ms [34] with different rates of horizontal transfer. If the gene transfer rate is zero, all 2803 genes evolve according to the same clonal genealogy of the population, i.e., one common species tree. In contrast, the individual gene trees may differ if some genes are effected by gene transfer. Nonetheless, the gene trees are still strongly dependent on each other due to the common link to the clonal genealogy. To investigate the effect of transfer on the accuracy of reconstruction, we used three different rates of gene conversion for the simulation of gene trees with ms (option −c with values 0, 2000, and 4000 with 6000 potential sites for gene conversion).

Among 2803 genes, 2100 are assigned to the most recent common ancestor (MRCA) at the root of the simulated gene tree while the remaining genes are gained at uniform distributed points at the branches after the MRCA on the gene tree. 300 of the 2100 ancestral genes are assigned to be present at all times, the remaining 2503 genes are lost at rate 2.1 along the branches of the corresponding gene tree as defined in [36] and [37]. After a loss event, the corresponding gene will be absent from all individuals descending from the branch of the loss event.

Given the gene trees and the presence absence pattern for each representative K-12 sequence, substitution can occur along the branches of the corresponding gene tree. We used seq-gen [38] to simulate these mutations according to the HKY model [39], setting the base frequencies to empirical *E. coli* base frequencies and the transition-transversion bias to 1.1. We simulated 5 sets of gene trees for no, occasional and frequent gene conversion. For each set we used 8 different substitution rate distributions to simulate pan-genomes: an exponential distribution with mean 0.06, uniform distributions between 0.05 and 0.1 and between 0.1 and 3, and constant substitution rates of 0.3, 0.2, 0.1, 0.05 and 0.01. The substitution rate *µ* of each gene was drawn from the corresponding distribution. The mean number of substitutions per site between two strains is given by 1*−e^−µT^*, where *T* is the distance between both strains in the gene tree.

## Results

### Benchmarking and comparison to other tools

To compare the clustering performance of different methods, one has to know which genes belong to the same cluster of orthologs. However, the orthoBench collection of manually curated groups of orthologous proteins [40] has been used to compare very diverged proteins across different domains of life and is far too diverse to benchmark a tool meant for pan-genomes of closely related bacteria. The orthologous groups in pan-genome datasets inferred using software tools depend on the methods used to generate these data sets and there is no ground truth. While pan-genomes based on real genomes can be used to compare clustering methods against each other, they are not immediately useful to assess accuracy.

To evaluate the performance of panX in an absolute sense and in comparison to state-of-the-art tools Roary [14], OrthoMCL [41], PanOCT [24], and OrthoFinder [42], we generated simulated pan-genomes for which the ground truth is known. In addition, we investigated the consistency of the orthologous clusters between tools in pan-genomes constructed from real bacterial genome sequences.

OrthoMCl and OrthoFinder were designed to identify orthologous groups across different domains of life, not for bacterial pan-genome inference. Nonetheless, they are often used in this context and we therefore included them here. In contrast, Roary was designed to cluster very large number of rather similar genomes. These tools are therefore expected to work well in different parameter ranges.

As a unique feature among all tools, panX relies on phylogeny based post-processing of the initial MCL clustering. This post-processing step is adaptive in that thresholds are scaled relative to the core genome diversity. As a result, panX works well across a large range of diversities.

#### Comparison of clustering accuracy on simulated datasets

We constructed pan-genomes by evolving 2803 genes from the E.coli genome along gene trees generated by the software ms [34]. Gene sequences evolve along the gene trees with different mutation rate distribution across the genome, can be gained or lost, and undergo horizontal transfer, see materials and methods for details.

We subjected 40 simulated pan-genomes of size 30 to analysis by Roary, PanOCT, OrthoFinder, OrthoMCL and panX and scored the resulting orthologous clusters. For each cluster, there are four possible outcomes: (i) A cluster is correct if it contains all and only genes from one true cluster, (ii) a cluster is incomplete but contains only genes from one true cluster, (iii) a cluster contains all genes from one true cluster but also genes from other clusters, or (iv) a cluster could fail on both counts. Fig. 2 shows how different tools perform at different levels of diversity of the pan-genome.

**FIG. 2:**
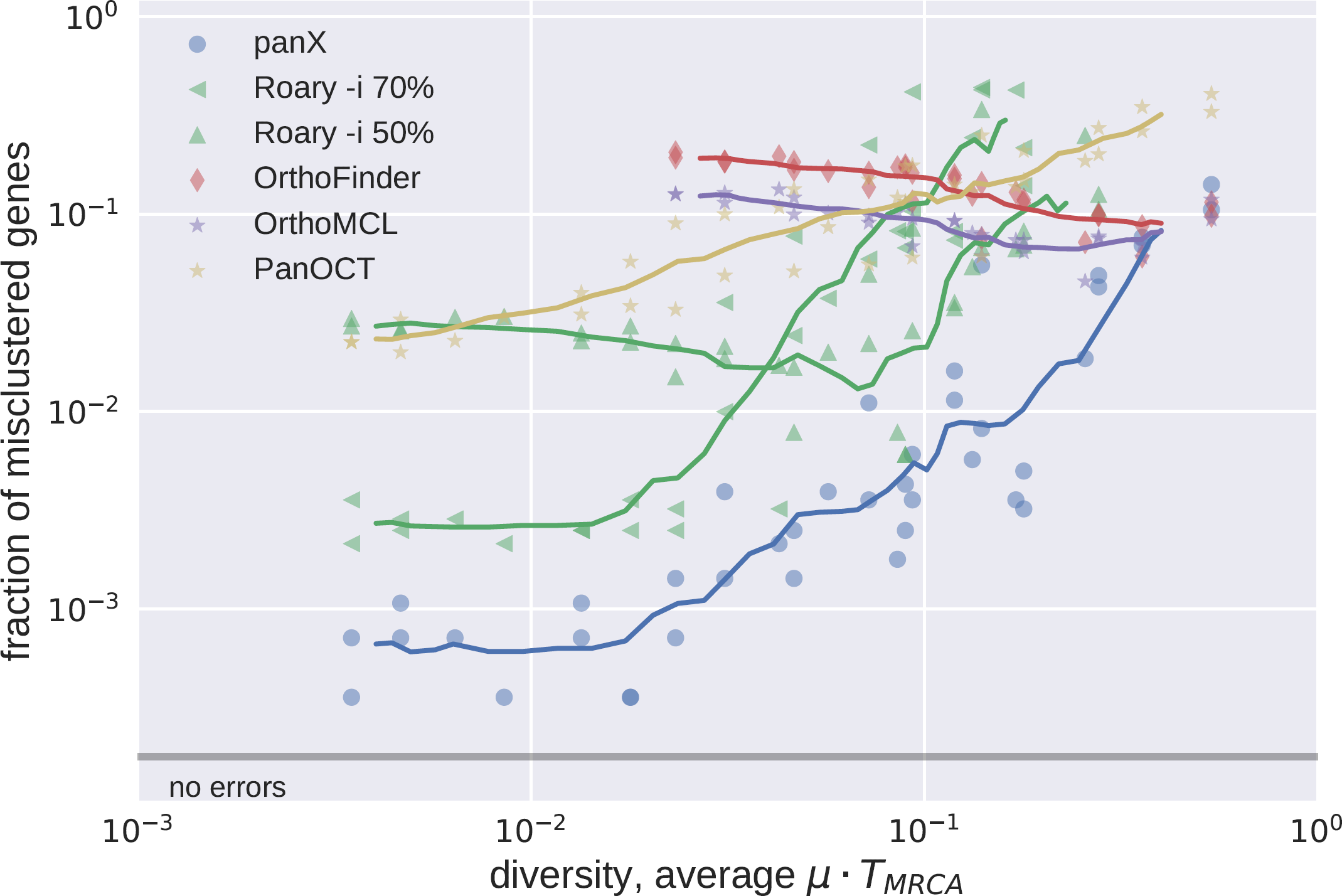
Accuracy of clustering by different tools. The fraction of misclustered genes increases with diversity of the pan-genome. We ran Roary with options −i 70 and −i 50. At low diversity, panX and Roary (-i 70) perform with similar accuracy and miscluster about 1 in 1000 genes. At high diversities, all tools have similar accuracy and miscluster 1 in 10 genes. Results for tools designed for high diversity data sets (OrthoMCL and OrthoFinder) are only shown for diversities above 0.02. Similarly, results for Roary are suppressed at high diversity to improve clarity of the graph.

OrthoMCL and OrthoFinder are designed for cross-species comparisons at large evolutionary distances. It is hence not surprising that these two tools merge many clusters that should be kept separate in low diversity pan-genomes. This effect is most pronounced for rare clusters, predominantly singletons, that get combined with other clusters, see Fig. 3. Core genes and other common gene clusters are typically correctly reconstructed. At very large diversities, OrthoMCL and OrthoFinder have an accuracy similar to that of panX. Roary and panX show similar behavior across a wide range of diversities from below 1% to 30% with panX typically making a factor of two fewer mistakes. However, we were unable to find a parameter set for Roary that worked well across the entire range of diversities. Using an identity cutoff of 70% (-i 70) worked best at low diversity, while lower cutoffs were required at high diversity. PanOct didn’t perform very well on our simulated data predominantly because it split too many clusters, see Fig. 3

**FIG. 3:**
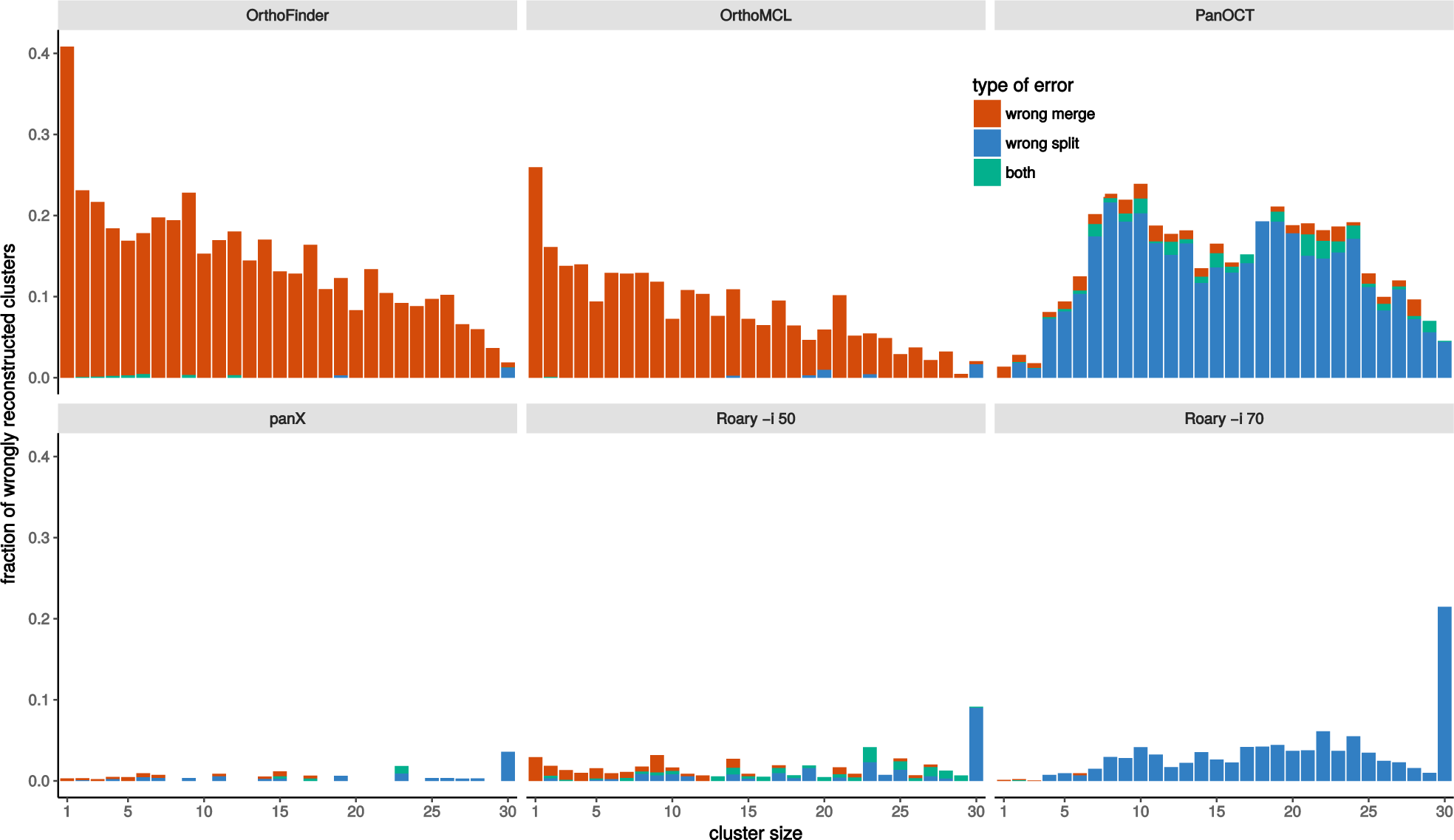
Type of misclustering by tool and gene frequency. The fraction of wrongly merged (red) and wrongly split (blue) clusters by gene frequency and clustering tool across 5 simulated datasets with exponentially distributed substitution rates with mean rate *µ* = 1/15.

The different types of errors (erroneous merging/splitting) are shown separately in Fig. S1. We repeated the analysis for different gene conversion rates and found mainly comparable results for no, occasional and frequent gene conversions (Fig. S2 and S3).

#### Real pan-genomes

While the ground truth of simulated pan-genomes is known, real pan-genomes lack an obvious point of comparison. Nonetheless, a comparison of the constructed clustering between different approaches can highlight the similarity and differences between them.

We used *S. pneumoniae* and *Prochlorococcus* pan-genomes to compare the results of the panX pan-genome identification pipeline to that of Roary, OrthoMCL, PanOCT, and OrthoFinder. We computed the size distribution of clusters, the size of the core genomes, and the total number of clusters from the pan-genomes estimated by different tools, see Fig. 4. For the low diversity collection of 33 *S. pneumoniae* genomes, the cluster size distributions inferred by the different tools are very similar (panel A) and the number inferred core genes differs by less than 10%. Between 78 and 86% of clusters inferred by panX are found by other tools. The greatest overlap is with Roary when run with identity threshold −i 70.

**FIG. 4:**
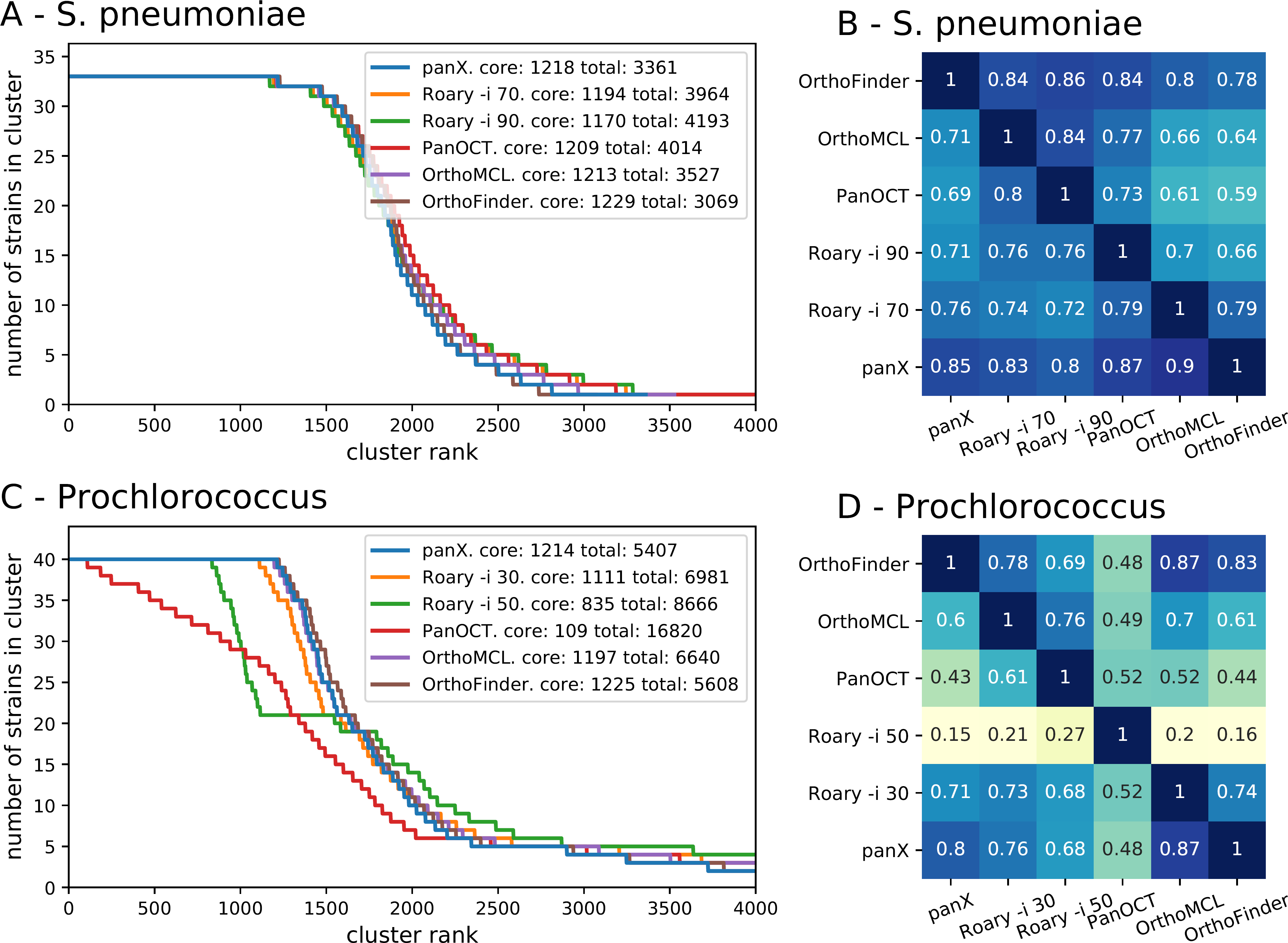
Pan-genome statistics: Panels A & C show the distribution of the number of strains represented in pan-genomes of 33 *S. pneumoniae* and 40 *Prochlorococcus* strains constructed by panX, Roary, OrthoFinder, OrthoMCL, and PanOCT (the last two tools are only available for the smaller *Prochlorococcus* data set). To obtain these graphs, clusters are sorted by descending number of strains represented in the cluster. This number is then plotted against the rank of the sorted clusters. The point where the lines drop below the number of strains marks the size of the core genome. PanX, OrthoFinder, and OrthoMCL largely agree on the cluster size distribution, the number of core genes and the total size of the pan-genome (with *∼* 10% variation). Roary agrees with the latter tools if the identity cut-off is chosen appropriately, while PanOCT estimates a very small core genome and an extremely large number gene clusters. Panels B & D show the degree to which different pan-genome tools agree with each other. Each row shows the fraction of clusters identified by one tool, that exactly match the a cluster identified by another tool. Analogous results for simulated data are given in Fig. S4.

More variation is observed in inferred pan-genomes of 40 *Prochlorococcus* strains, see Fig. 4C&D. Roary [14] separates nearly all *Prochlorococcus* genes and identifies only 10 core genes when using standard parameters. After lowering the minimum percentage identity for blastp in Roary to 30% (-i 0.3), Roary identified 1111 core genes vs 1214 identified by panX. While Roary warns that it has not been designed to support such diverse datasets, the 60% of the resulting clusters agree with those identified by panX. PanX and Roary identified 5,407 and 6,981 clusters of orthologous genes, respectively – not too far from the estimated pan-genome size of *>* 8,500 genes present in more than one percent of the population [43]. Although each tool constructs a number of unique clusters, the results for OrthoFinder and OrthoMCL are comparable to those of Roary and panX. In contrast, the tool PanOCT separates many more genes than panX or Roary with parameter −i 30. PanOCT is designed for closely related prokaryotic strains and therefore splits the diverse *Prochlorococcus* genomes into 16,820 clusters with mainly 1-3 genes. Only 109 core clusters are identified. The tools OrthoMCl and OrthoFinder, designed for more diverse data sets, generate cluster size distributions similar to those by panX and between 71 and 87% of clusters found by one tool are also found by another.

Testing this collections of pan-genome tools on larger data sets proved prohibitive since only Roary and panX scale well with number of genomes.

### Web application for pan-genome exploration

To explore the pan-genome constructed by the pipeline described above, we developed a browser based visualization. The layout of the application is that of a large dashboard (see Fig. 5), on which multiple aspects of the pan-genome can be interrogated simultaneously.

**FIG. 5:**
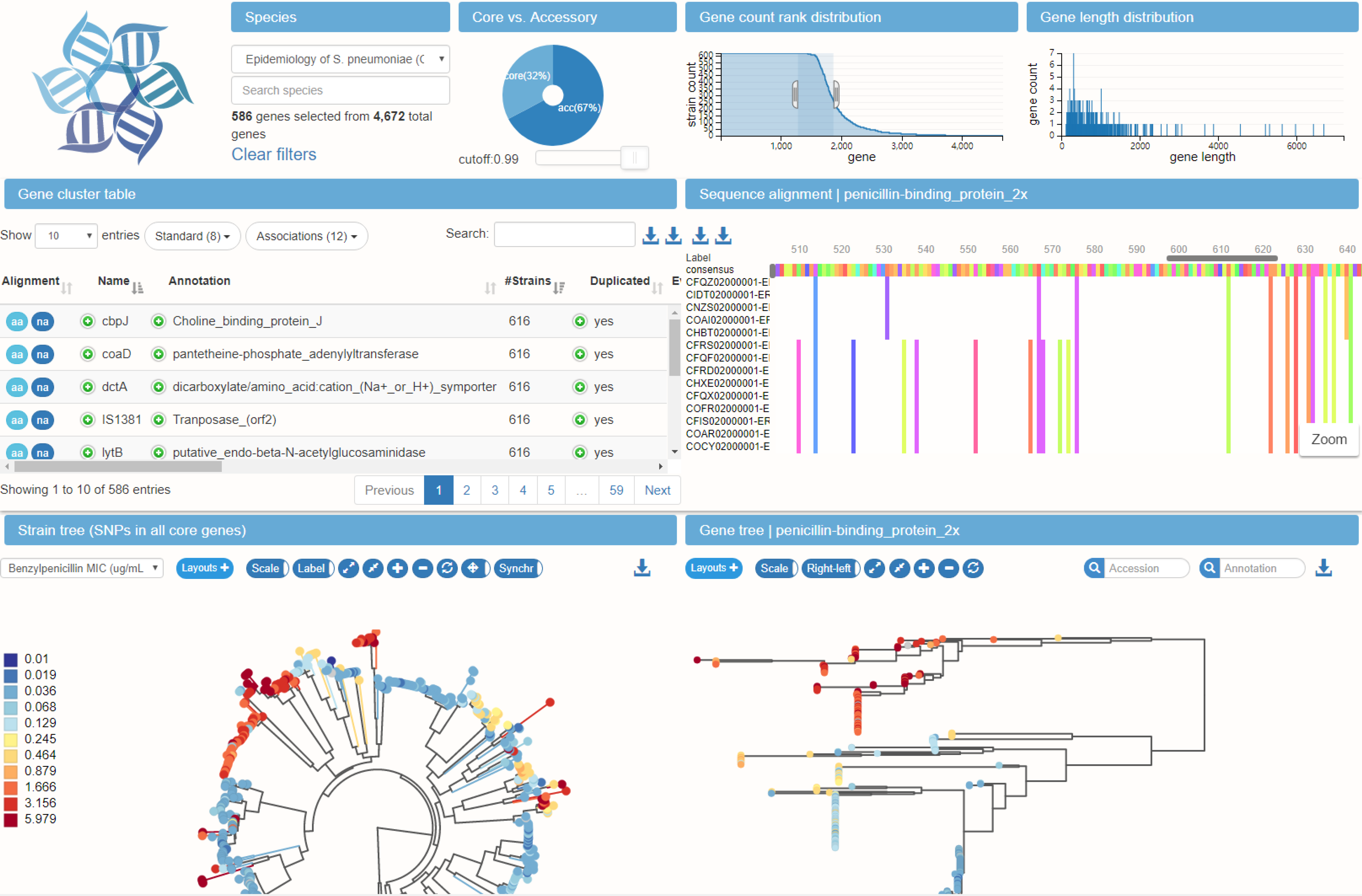
Interconnected components of the panX web application: The top panels provide a statistical characterization of the pan-genome and allow filtering of gene clusters by abundance and gene length. The gene cluster table on the center left is searchable and sortable and allows the user to select individual gene clusters for closer inspection. Upon selection in the table, the alignment of gene cluster is loaded into the viewer on the center right, the gene tree is loaded into the tree viewer at the bottom right, and presence/absence patterns of this gene cluster are mapped onto the core genome tree at the bottom left. The example shows the gene coding for the penicillin binding protein Pbp2x and the color indicates the MIC against benzylpenicillin.

At the top, three graphs provide basic statistics on the abundance and length distribution of all genes. In the middle row, a searchable table contains summary statistics and annotations for all gene clusters. The alignment viewer on the right shows the nucleotide or amino acid alignment of gene cluster selected in the table. Below the table, the core genome SNP tree is shown, along with a phylogenetic tree of the currently selected gene cluster. At the very bottom, a second searchable table allows rapid access to meta information available for different strains.

The hallmark of the panX web-application are the interconnected components that illustrate different properties of the gene clusters. The pan-genome statistic charts at the top allow rapid sub-setting of gene clusters by gene length and abundance. The left chart shows an inverse cumulative distribution of clusters sizes, i.e., clusters are sorted by decreasing number of strains represented in the cluster, such that all core genes present in all strains are shown on the left. The size of the core genes is then simply the length of the plateau of the curve to the first drop. The core genome is followed by gradual decline in gene number from common to rare accessory genes. Lastly, a long tail contains the strain-specific singletons. Subsets of genes can be easily defined by selecting a range of the graph with the mouse. Similarly, the center chart shows the distribution of gene length.

The pie chart on the right shows the proportion of core and accessory genome, each of which can be selected by clicking on the sectors in the chart. To allow for soft and strict definitions of the core genome, the cut-off delineating core and accessory genome can be adjusted with a slider.

#### Rapid and searchable access to alignment and gene trees

The table of all gene clusters is dynamically restricted to the range of gene abundances and gene lengths selected above. The table can be searched by gene name and annotation or sorted by gene count, diversity etc. Annotations of all input sequences (also discordant annotations of genes belonging to the same gene cluster) are accessible by expanding the annotation field. Similarly, the column *duplicated* specifies whether the gene cluster contains more than one gene per strain. The list of strains in which genes are duplicated and copy number of this gene can be accessed by expanding the row. Each row contains triggers to show the corresponding nucleotide or amino acid sequence alignment in the alignment viewer (MSA) from BioJs [44]. In order to high-light difference among sequences, only consensus sequence and variable sites are shown by default, while the corresponding original alignment can be downloaded. This trigger also updates the phylogenetic tree viewers. Searching *mcr-1*, for example, immediately highlights the 11 *E. coli* genomes in the RefSeq database that have an annotated mobile colistin resistance gene.

#### Interactive core genome tree and gene tree viewers

To facilitate the comparison between the core genome SNP tree and the gene tree, the two trees have connected interactive elements. When placing the mouse on a leaf node in one tree, the corresponding nodes are highlighted in both trees. Similarly, if the mouse is placed over an internal node, all nodes in the corresponding clades are highlighted with different colors for each strain. This gives a rapid impression of whether the core genome tree and the gene tree are compatible and whether the gene is duplicated in some of the strains, see Fig. 6.

**FIG. 6:**
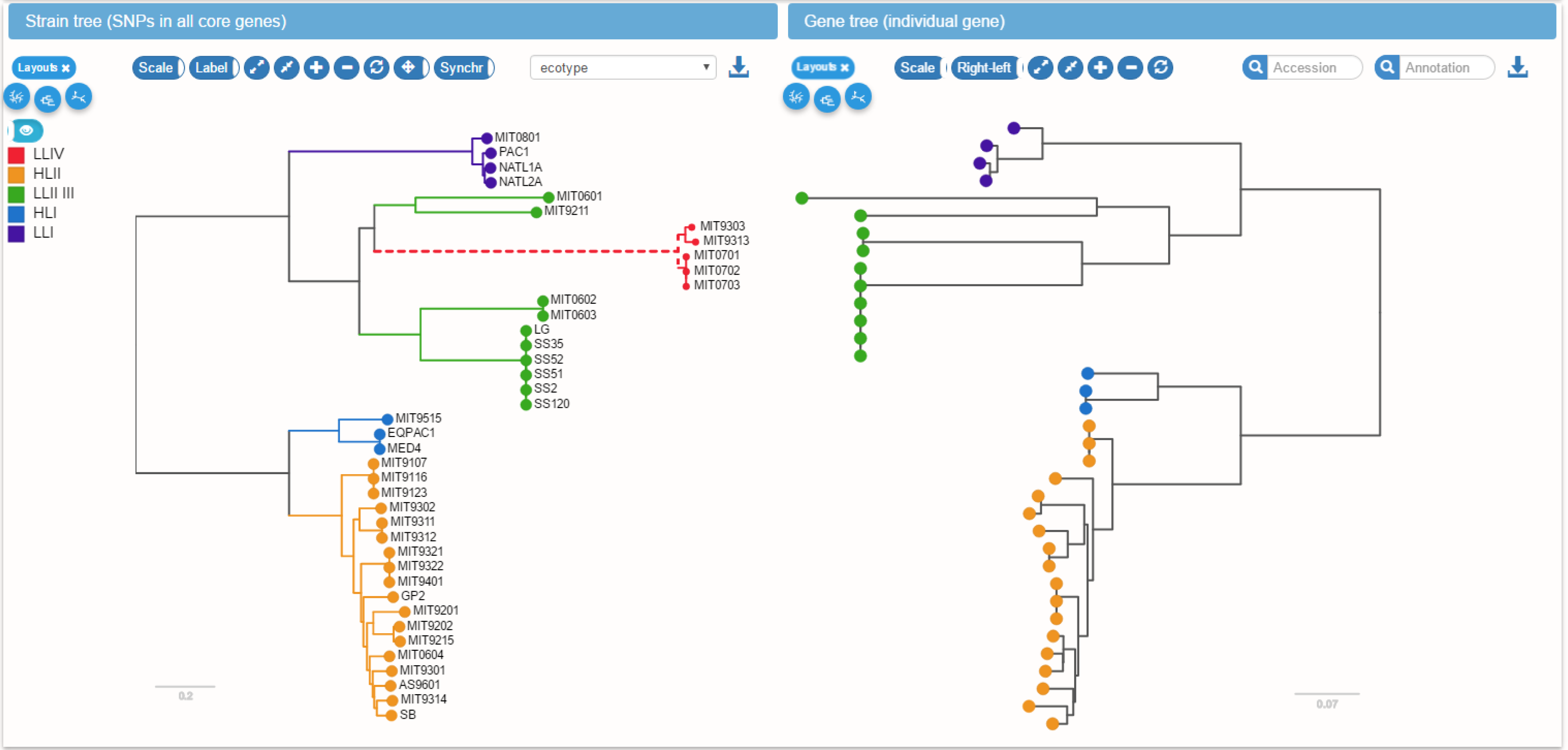
Linked core genome and gene trees. The core genome tree shows the strains in which the current gene is present or absent. Placing the mouse over an internal node in one of the trees (upper clade of the gene tree on the right in this example) highlights all strains in the corresponding clade in both trees. This gives the user a rapid impression of phylogenetic incongruence and likely gene gain and loss events.

The most likely gene loss and gain events inferred by the ancestral reconstruction algorithm are indicated on the tree by dashed or thick lines, respectively. Mutations in the amino acid or nucleotide sequence of the gene are mapped onto the gene tree and can be inspected using the tooltips associated with branches in the tree.

In addition to mutation and gain/loss events, the tree can be colored with metadata associated with different strains. Such metadata would typically include collection dates, sampling location, host species or resistance phenotypes.

### Pan-genomes of common bacterial groups

We ran the panX on collections of genomes of bacterial groups for which more than 10 genomes were available in RefSeq resulting in a total of 94 pan-genomes including many human pathogens. Statistics of a subset are shown in Tab. I, the corresponding data for all species are given as supplementary material. The majority of these species exhibit low diversity in their core genomes with typically just a few percent nucleotide differences, some times even less than 0.001. The median number of core genes is 1800 while the median pangenome size is about 5000 genes. Diverse species tend to have smaller core-genomes and larger pan-genomes, as expected.

### Pan-genomes of diverse collections of genomes

Most of these collections are closely related genomes, but we also included a diverse group of genomes of *Prochlorococcus* and the pan-genomes of the bacterial orders Pseudomonadales, Enterobacteriales and Vibrionales.

*Prochlorococcus* is a marine cyanobacterium that is responsible for a significant fraction of the marine primary production and serves as a model system in marine microbial ecology [45]. While we relied on annotations available in NCBI for most species, we re-annotated the genomes of 40 *Prochlorococcus* sequences [46] using Prokka [47]. The annotation was derived from a custom database based on the 12 annotated *Prochlorococcus* strains CCMP1375, MED4, MIT9313, NATL2A, MIT9312, AS9601, MIT9515, NATL1A, MIT9303, MIT9301, MIT9215 and MIT9211. *Prochlorococcus* is a much more diverse population than the other species we investigated, see Tab. I, which makes it a challenging case for pan-genome analysis.

While the 16S rRNA sequences of all 40 *Prochlorococcus* strains do not differ by more than 3%, *Prochlorococcus* can be divided into ecotypes that are remarkably different in genome size and GC content [45]. These ecotypes correspond to high and low light intensity adapted *Prochlorococcus* populations and can be visualized along the species and gene trees in panX. While the genomes of *Prochlorococcus* have likely been streamlined by strong selective forces to lose genes [48], gene gains and duplications have frequently occurred in all *Prochlorococcus* lineages. Two well-known examples for *Prochlorococcus* are the gain of nitrate assimilation genes *nirA* [49] and gene duplication and phage mediated gene transfer of the photosynthesis gene *psbA* [50]. The ancestral insertions of these genes are placed on the species tree by panX and can be investigated in the strain phylogeny if one searches for the gene name and gene presence/absence is chosen as metadata.

In addition to the diverse *Prochlorococcus* genomes, we analyzed collections of genomes that encompass the entire bacterial orders *Pseudomondales*, *Enterobacteriales*, and *Vibrionales*. For each of these orders, we collected at most 10 genomes from each species (based on the species designation in the RefSeq files) to avoid over-representation of human pathogens. Running panX on these orders resulted in about 1000 core genes for *Pseudomondales* and *Vibrionales* and about 2000 core genes in case of the smaller collection of *Enterobacteriales*. Core genes of *Pseudomondales* and *Vibrionales* were typically 20% diverged from each other, while *Enterobacteriales* core genome was less diverse at 11%. The core genome SNP tree of the *Vibrionales* clearly separates the genomes by species, while the core genome trees of the other orders show considerable mixing of species.

### Large pan-genome of *Streptococcus pneumoniae*

The utility of the interactive web application is most evident for collections of genomes with rich meta data. One such collection is the *S. pneumoniae* data set generated by **(author?)** [51]. This data set consists of 616 whole genome sequences and rich meta data including antibiotic susceptibility and host characteristics. Even with 616 genomes, the web application is fluid and responsive.

For S. pneumoniae, we calculated branch and presence/absence association scores for every gene cluster. All scores are included in the gene cluster table, which can be sorted by each score. While many strongly associated genes are false positive, they are enriched for known genes. The gene cluster with the largest branch association with benzylpenicillin MIC is the penicillin binding protein *pbp2x*. The coloring of the gene tree by the benzylpenicillin MIC confirms that the resistant and susceptible isolates form two distinct clades separated by a large number of amino acid substitutions, see Fig. 5. While resistant strains are scattered across the species tree, they form a single clade in the tree of *pbp2x*.

Similarly, the gene cluster table can be sorted by presence/absence association scores. The gene cluster that is most associated with erythromycin resistance is *mefE* coding for an efflux pump.

### Availability

The computational pipeline to identify the pan-genome consists of a collection of python scripts and a master script that runs desired analysis steps in series. The visualization is build on node.js server and makes extensive use of BioJS [44], D3.js [52], `dc.js` [53], and other javascript libraries. The analysis pipeline and the code for the web application is made available under the GPL3 license on [github](https://github.com/neherlab) as repositories *pan-genome-analysis* and *pan-genome-visualization*.

The web application can either be hosted on a web server or can be used locally to inspect and explore pan-genomes produced by the panX pipeline. We computed a large number of pan-genomes and made those available at pangenome.de. The website currently hosts 93 bacterial species including those listed in Tab. I. Several downloading options are available: core gene alignments and all gene alignments can be down-loaded via the down-arrow button next to the gene cluster table. Alignments and gene trees for individual clusters and the strain tree can be downloaded via buttons next to the alignment viewer and tree viewers, respectively. All buttons are associated with tooltips explaining the action of the buttons.

**TABLE I:** Summary statistics of pan-genomes available at pangenome.de. Average number of pairwise differences per nucleotide in core gene alignments.

| Species | genomes | core genes | all genes | singletons | diversity* |
| --- | --- | --- | --- | --- | --- |
| *Acinetobacter baumannii* | 71 | 1701 | 8334 | 1558 | 0.010 |
| *Bacillus anthracis* | 43 | 4156 | 5980 | 62 | 1.0e-04 |
| *Bacillus cereus* | 36 | 2979 | 13364 | 3486 | 0.048 |
| *Bordetella pertussis* | 291 | 2437 | 3743 | 158 | 4.1e-06 |
| *Burkholderia pseudomallei* | 59 | 4098 | 11580 | 1966 | 0.003 |
| *Campylobacter jejuni* | 113 | 935 | 3166 | 526 | 0.014 |
| *Chlamydia trachomatis* | 68 | 809 | 978 | 12 | 0.005 |
| *Clostridium botulinum* | 23 | 795 | 9083 | 2294 | 0.147 |
| *Corynebacterium pseudotuberculosis* | 59 | 1133 | 2316 | 65 | 0.005 |
| *Enterobacter cloacae* | 22 | 2971 | 10783 | 3211 | 0.087 |
| *Escherichia coli* | 307 | 778 | 23107 | 6339 | 0.015 |
| *Francisella tularensis* | 35 | 838 | 2339 | 302 | 0.007 |
| *Helicobacter pylori* | 85 | 694 | 2371 | 328 | 0.042 |
| *Klebsiella pneumoniae* | 109 | 2545 | 15978 | 4004 | 0.007 |
| *Listeria monocytogenes* | 95 | 1907 | 4947 | 485 | 0.031 |
| *Mycobacterium tuberculosis* | 51 | 2665 | 4350 | 93 | 2.0e-04 |
| *Neisseria meningitidis* | 78 | 1071 | 3375 | 426 | 0.015 |
| *Prochlorococcus marinus* | 40 | 1047 | 5407 | 1262 | 0.291 |
| *Pseudomonas aeruginosa* | 70 | 3264 | 12768 | 3195 | 0.006 |
| *Salmonella enterica* | 260 | 1327 | 15521 | 3996 | 0.009 |
| *Staphylococcus aureus* | 146 | 1229 | 5206 | 731 | 0.008 |
| *Streptococcus pneumoniae* | 33 | 1188 | 3361 | 540 | 0.010 |
| *Streptococcus pyogenes* | 50 | 970 | 2856 | 341 | 0.008 |
| *Vibrio cholerae* | 28 | 2412 | 5156 | 771 | 0.005 |
| *Xanthomonas citri* | 26 | 3385 | 5261 | 291 | 0.001 |
| *Yersinia pestis* | 33 | 2557 | 4587 | 172 | 1.0e-04 |
| Pseudomonadales | 119 | 966 | 42520 | 20577 | 0.194 |
| Enterobacteriales | 33 | 1998 | 16413 | 6988 | 0.112 |
| Vibrionales | 66 | 716 | 30461 | 15643 | 0.193 |

## Conclusions

Being able to visualize and explore high dimensional data is often the key to developing insight into the mechanisms driving complex dynamics. PanX is meant to enable such exploration of large sets of bacterial genomes, which are characterized by the evolution of individual genes as well as the gain and loss of genes. The design of panX focused on combined breadth and depth: Besides summary statistics and species trees, panX allows to select interesting sets of genes or search for individual genes. Alignments and phylogenetic trees of genes can then be analyzed in detail with individual mutations and gain/loss events mapped to the gene tree and the core tree, respectively. The evolutionary patterns of genes can then be compared to meta-information such as resistance phenotypes associated with the individual strains.

By integrating meta-information with the molecular evolution of genes and genomes in one visualization, panX can assist investigations of the dynamics of pan-genomes and adaptation of bacteria to new habitats and environmental challenges. Horizontal transfer is pivotal for many aspects of bacterial adaptation [11], but at the same time, it remains much more difficult to analyze than evolution by vertical descent [3]. The ability to interactively explore such pan-genomes might help to grasp the complexity of this dynamics.

On the other hand, a web-based tool that can be readily kept up-to-date by addition of newly sequenced isolates would be useful in pathogen surveillance. When paired with meta-information such as resistance, pathogenicity, sampling date, location and comorbidities, panX can help to study adaptation, spread, and transmission chains of pathogens. Similar approaches have proved useful at tracking spread and evolution of seasonal influenza virus or Ebola virus during the recent outbreak in West Africa [29, 54]. Currently, phenotype data are available for a minority of the whole genomes sequences and data sets like the *S. pneumoniae* by (author?) [51] are an exception. With increasing availability and timely publication of such data from routine surveillance, panX or derivatives could be used to track foodborne outbreaks, monitor the global spread of drug resistance bacteria [55], or assist infection control in individual hospitals. The time required to build a pan-genome of 1000 strains is less than a day on a 64 core node such that frequent updates of such a tracking tool are possible.

## Acknowledgments

We gratefully acknowledge stimulating discussions with Matthias Willmann and Erik van Nimwegen as well as advice on `dc.js` from Gordon Woodhull.

